# Medaka population genome structure and demographic history described via genotyping-by-sequencing

**DOI:** 10.1101/233411

**Authors:** Takafumi Katsumura, Shoji Oda, Mitani Hiroshi, Hiroki Oota

**Author notes:** **Authors for Correspondence:** Takafumi Katsumura, Graduate School of Natural Science and Technology, Okayama, University, Okayama, Japan, Hiroki Oota, Department of Anatomy, Kitasato University School of Medicine, Kanagawa, Japan.

## Abstract

Medaka is a model organism in medicine, genetics, developmental biology and population genetics. Lab stocks composed of more than 100 local wild populations are available for research in these fields. Thus, medaka represents a potentially excellent bioresource for screening disease-risk- and adaptation-related genes in genome-wide association studies. Although the genetic population structure should be known before performing such an analysis, a comprehensive study on the genome-wide diversity of wild medaka populations has not been performed. Here, we performed genotyping-by-sequencing (GBS) for 81 and 12 medakas captured from a bioresource and the wild, respectively. Based on the GBS data, we evaluated the genetic population structure and estimated the demographic parameters using an approximate Bayesian computation (ABC) framework. The autosomal data confirmed that there were substantial differences between local populations and supported our previously proposed hypothesis on medaka dispersal based on mitochondrial genome (mtDNA) data. A new finding was that a local group that was thought to be a hybrid between the northern and the southern Japanese groups was actually a sister group of the northern Japanese group. Thus, this paper presents the first population-genomic study of medaka and reveals its population structure and history based on autosomal diversity.

## Introduction

Medaka (*Oryzias latipes*) is a small fresh-water fish native to East Asia that has attracted attention as a vertebrate model for population genetics (Oota and Mitani 2011; Spivakov *et al*. 2014). Wild medaka populations have been maintained in certain universities and research institutes as a bioresource (hereafter, wild lab stocks) with funding from the Japanese government since 1985 (Shima *et al*. 1985). These populations consist of more than 100 local-wild populations that have various phenotypic traits (Watanabe-Asaka *et al*. 2014; Igarashi *et al*. 2017) and abundant genetic diversity (Kasahara *et al*. 2007). Geographical features affect the population structure of organisms. Seas and mountains restrict the movement of terrestrial animals and freshwater fish, and the climate also changes the breeding timing and foraging environment. Particularly, noticeable climate differences are observed within island groups, such as the Japanese archipelago, where a large latitudinal difference exists between the southern and northern ends. Therefore, animals are exposed to various selective pressures according to the geographic environment. The local populations have differentiated into local groups by genetic drift, resulting in genetically divergent groups. Medakas stocked as a bioresource are thought to have retained the genetically adapted traits they acquired from various environments of the Japanese archipelago.

Our ultimate goal in exploiting medaka characteristics is to establish an experimental system for testing the functional differences between alleles detected by, for instance, genome-wide association studies. Once genetic polymorphisms related to phenotypic traits in medaka are detected, *in vivo* experiments, such as genome editing experiments (Ansai and Kinoshita 2014), can be conducted to understand the functions of the genetic variants (Shimmura *et al*. 2018). Revealing the functional difference between alleles in wild populations allows us to infer the role of genetic polymorphisms in human homologous genes (Igarashi *et al*. 2017; Shimmura *et al*. 2018).

Most previous analyses of medaka genetic diversity and population structure have been conducted using mitochondrial DNA (mtDNA). Medaka is divided into four mitochondrial groups: the northern Japanese (N.JPN), southern Japanese (S.JPN), eastern Korean (E.KOR) and western Korean/Chinese groups (W.KOR) (Sakaizumi *et al*. 1980; 1983; Sakaizumi 1986; Katsumura *et al*. 2009). These groups do not have sympatric habitats. Although previous allozyme-based studies show that an ambiguous group exists in the geographic boundary region known as Tajima-Tango (see also fig. 1, supplementary fig 1), between N.JPN and S.JPN, which is thought to be a hybridization of the two groups (Sakaizumi 1984; Takehana *et al*. 2016), the process has not been fully verified.

**FIG. 1:**
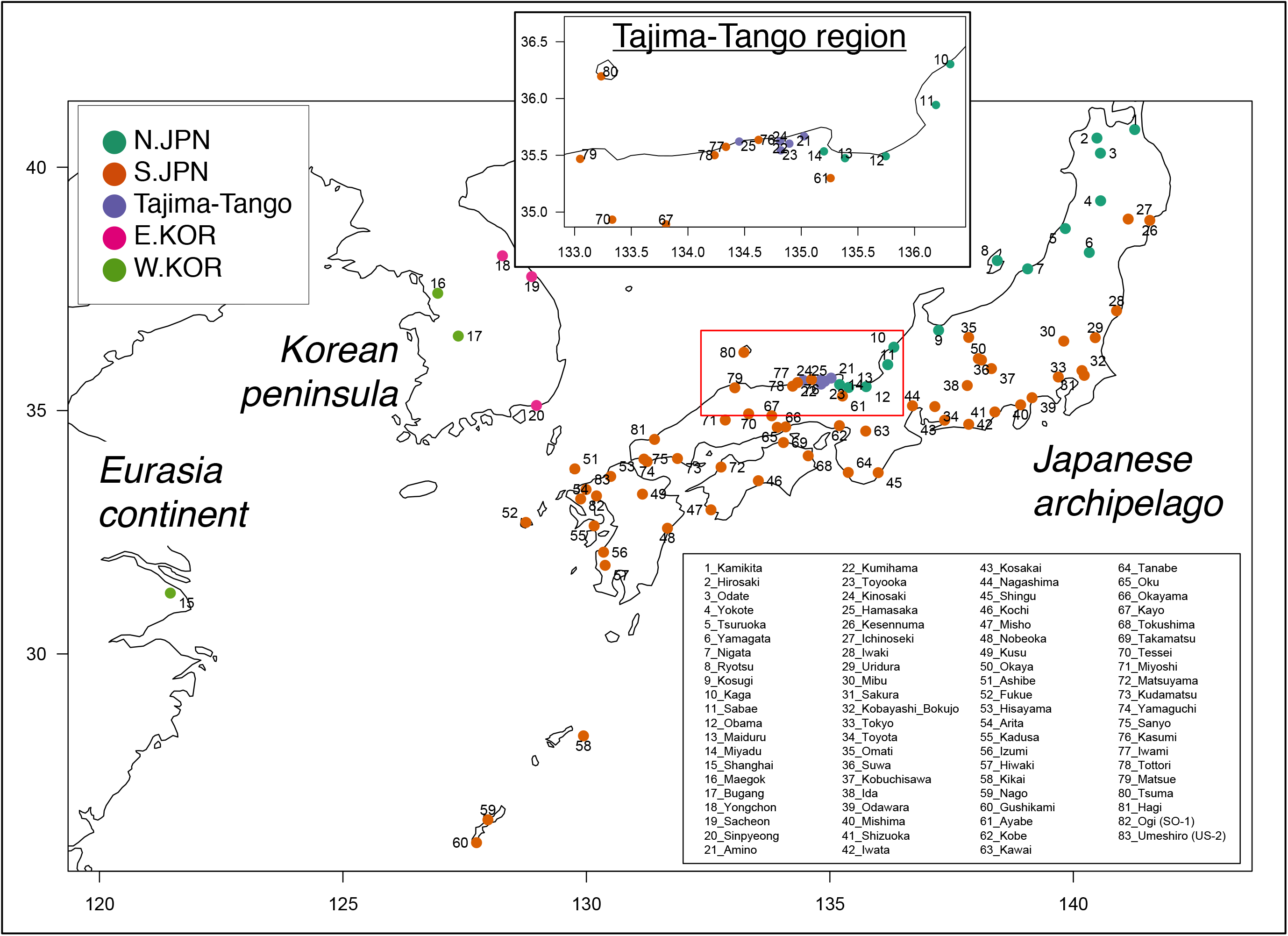
Map of the original locations of the wild lab stocks and wild-captured medakas. In the upper central map, the enlarged red frame shows the boundary region between S.JPN and N.JPN. Each color represents the mtDNA and allozyme-based groups shown in the left upper inset box. The numbers on the map are consistent with the population IDs, with the names on the right bottom inset box.

Based on mtDNA cytochrome *b* gene sequences, N.JPN and S.JPN are composed of 3 and 11 subgroups, respectively (Takehana *et al*. 2003). Each subgroup is composed of local populations, and the between-population genetic diversities are greater than the within-population genetic diversities, indicating substantial genetic differentiation between local populations (Katsumura *et al*. 2009). Their habitat environments are also largely different. For instance, there is a large amount of snowfall in the habitats of N.JPN, where the breeding season is short. The habitat of S.JPN is wider than that of N.JPN, and its climate environment is also diverse; e.g., the differences of the annual average temperature and rainfall between the south end (Nago in Okinawa) and north end (Ichinoseki in Iwate) inhabited by S.JPN were 11.6 °C and 988 mm in 2017, respectively (http://www.jma.go.jp/jma/indexe.html). The phylogenetic data from these studies also suggested that N.JPN and S.JPN have been spreading in the Japanese archipelago at different times. Particularly, the origin of S.JPN, which has the largest habitat, has been suggested to be the northern part of Kyushu Island based on mtDNA (“Out of Northern Kyushu” hypothesis) (Katsumura *et al*. 2012). However, mtDNA is insufficient to describe the population structure and history because of its single locus. Genetic diversity based on autosomes is essential to understand the medaka population structure, but the comprehensive data of autosomes are still limited.

An inference of the population structure and history induced by autosomal information is more robust than that induced by single loci. To unravel the population structure based on autosomes, we comprehensively performed a population-genetic analysis based on autosomal single-nucleotide polymorphisms (SNPs) using all 81 local populations maintained as wild lab stocks at the University of Tokyo. We examined an individual sampled from each population stock and 12 wild individuals captured from the northern part of Kyushu Island, where (Katsumura *et al*. 2012) the medakas currently distributed along the entire Pacific side of the Japanese archipelago originated, to estimate each population’s time of expansion from Northern Kyushu. To obtain SNP data, we conducted genotyping-by-sequencing (GBS) (Elshire *et al*. 2011; Narum *et al*. 2013) using a high-throughput sequencer. This method, which allowed us to cost-effectively genotype tens of thousands of SNPs (Andrews *et al*. 2016), resulted in an accurate enhancement of the population-genetic estimations because the variance in the demographic parameter estimates decreased when we used many SNPs. Eventually, we obtained more than ten thousand SNPs from eighty-one wild lab stocks and twelve wild-captured medakas. Here, we re-evaluated the medaka genetic diversity and population structures based on these SNPs and reconstructed the population history by assessing three demographic events. These data redefine the medaka local groups and provide a basis of the population history for discussing the role of phenotype-associated alleles in the context of adaptation.

## Materials and Methods

### Samples

We sampled 81 male medakas from 81 wild lab stocks (14 from N.JPN, 56 from S.JPN, 5 from Tajima-Tango in the geographic boundary region between N.JPN and S.JPN, 3 from E.KOR, and 3 from W.KOR; note that these groupings come from previous mtDNA sequences and allozyme patterns) at the University of Tokyo in 2014. The lab stocks originated from geographically distinct populations in East Asia and have been maintained since 1985 (Shima *et al*. 1985) as closed colonies in the Graduate School of Frontier Sciences, the University of Tokyo, Kashiwa City (fig. 1). These lab stocks maintain the genetic diversity originating from their habitat (Katsumura *et al*. 2014; Igarashi *et al*. 2017) and show less diversity within populations than between populations (Katsumura *et al*. 2009).

Because of these characteristics, we considered that one individual sampled from the lab stock would be adequate to represent its originating population. In addition, we used two wild-captured medaka populations from the Saga Prefecture in the northern part of Kyushu, Japan. One population was the Ogi (SO) population from Southern Saga, and the other population was the Umeshiro (US) population from Northern Saga, both of which were captured in September 2010 (see Katsumura *et al*. 2012). Six of 48 medakas from each wild-captured population were selected randomly and analyzed via GBS.

### DNA extraction and genotyping-by-sequencing

One-third of the medaka body was dissolved in a 600 μl lysis buffer containing 1.24% SDS, 0.124 M EDTA and 0.062 mg/ml proteinase K (final concentrations). The total genomic DNA was extracted and purified using phenol-chloroform and isopropanol precipitation.

After a 70% EtOH wash, an isolated DNA pellet was resuspended in 100 μl TE buffer and then treated with RNase A (final conc. 1 mg/ml) for 1 hour at room temperature. Then, the DNA was purified again using phenol-chloroform and isopropanol precipitation. For ninety-three samples, the GBS process was outsourced to Macrogen Japan in Kyoto. The procedures for constructing libraries and performing Illumina HiSeq 2000 single-end sequencing were the same as those described by Poland *et al*. (Poland *et al*. 2012) except for the use of the restriction enzyme ApeKI instead of EcoRI–Mspl. The sequence lengths were 51 bp and included each individual in-line barcode (4–9 bp) for the individual sample. The data have been submitted to the DDBJ Sequence Read Archive (DRA) database under project accession ID: DRA006353.

### Quality filtering and SNP extraction

Our single-end reads were filtered using *FASTQ Quality Filter* in *FASTX-Toolkit version 0.0.13* (http://hannonlab.cshl.edu/fastx-toolkit) using the following options: -Q 33 -v -z -q 30 -p 90. The draft genome of the medaka sequenced by the PacBio sequencer (Medaka-Hd-rR-pacbio_version2.0.fasta; http://utgenome.org/medaka_v2/#!Assembly.md) was used to align the reads using *BWA backtrack 0.7.12-r1039* (Li and Durbin 2009) using the “-n 0.06” option. After the mapping process, the multi-mapped reads were removed using *Samtools v1.2* (Li *et al*. 2009) and the “-Sq 20” option. Following this pipeline, sequencing of the ApeKI-digested GBS libraries generated an average of 3.06 million reads per individual before any quality filtering. The read numbers ranged from 1.89 to 4.10 million reads per individual. After quality filtering, 2.76 million (90.2%) sequences per individual on average were retained, and 0.30 million (9.8%) sequences were eliminated. The retained sequences presented a mean quality score of 38.7 and a GC content of 47.8%. An average of 1.84 million of the retained sequences (66.7%) aligned to the medaka autosomal genome, and 0.92 million sequences (33.3%) were not aligned and discarded because they mapped to non-autosomal loci (mitochondrial genome and unanchored contigs) and multiple loci.

The *Stacks* pipeline (version 1.35) and the Stacks workflow (https://github.com/enormandeau/stacks_workflow) were used to generate SNPs and sequences for each individual separately (Catchen *et al*. 2011; 2013). The selected *Stacks* parameters were as follows: minimum stack depth (-m), 3; and number of mismatches when building the catalog (-n), 1. In the ‘rx’ step of *Stacks*, we used the bounded SNP model set to 0.1, the ε upper bound set to 0.1 and the log likelihood set to 10. Following this pipeline, five datasets were constructed using the *population* program in *Stacks*.

The first was the “PopStat” dataset, which was generated using the “-p 5 -r 0.66” options and mitochondrial grouping (N.JPN, Tajima-Tango, S.JPN, E.KOR, W.KOR; see also fig. 1) to calculate the population-genetic statistics (table 1) using the loci shared across all groups and sequenced in one or more populations in each group.

**Table 1.** Summary genetic statistics for five populations using “PopStat” dataset. These statistics include the mean number of individuals genotyped at each locus (*1), the number of variable sites in five populations (Variants), the number of polymorphic sites in each population (Polymorphic) and the number of variable sites unique to each population (Private). The number in parenthesis is Standard Error.

| Group | Number of Individual*<sup>1</sup> | Total length of GBS loci (bp) | Variant | Polymorphic | Private | Major Allele Frequency | Observed Heterozygosity | Nucleotide Diversity |
| --- | --- | --- | --- | --- | --- | --- | --- | --- |
| N.JPN (SJ.NEH) | 11.5<br>(+/-0.03) | 45968 | 2453 | 256 | 136 | 0.983<br>(+/-0.0014) | 0.023<br>(+/-0.0022) | 0.0014<br>(+/-0.0001) |
| Tajima -Tango | 4.4<br>(+/-0.01) | 44011 | 2453 | 199 | 71 | 0.980<br>(+/-0.0016) | 0.024<br>(+/-0.0022) | 0.0017<br>(+/-0.0001) |
| S.JPN | 41.6<br>(+/-0.08) | 45968 | 2453 | 1448 | 1231 | 0.958<br>(+/-0.0017) | 0.029<br>(+/-0.0016) | 0.0036<br>(+/-0.0001) |
| W.KOR | 2.4<br>(+/-0.01) | 44990 | 2453 | 467 | 342 | 0.940<br>(+/-0.0027) | 0.054<br>(+/-0.0032) | 0.0053<br>(+/-0.0003) |
| E.KOR | 2.4<br>(+/-0.01) | 44011 | 2453 | 346 | 203 | 0.953<br>(+/-0.0025) | 0.041<br>(+/-0.0030) | 0.0042<br>(+/-0.0002) |

The second was the “Global” dataset, which was generated using the “-p 58 -r 1.00” option and without the mitochondrial grouping, to examine the phylogenetic relationships between the geographic populations and the population structure within the species.

The third and the fourth were the “HZ-1” and “HZ-2” datasets (“HZ” is the abbreviation of “Hybrid zone”), respectively, including the 15 boundary populations (N.JPN: Kaga, Maiduru, Miyadu, Obama, and Sabae; Tajima-Tango: Amino, Hamasaka, Kinosaki, Kumihama, and Toyooka; and Honshu: Ayabe, Iwami, Kasumi, Matsue, and Tottori) with or without the Kyushu populations (Kyushu: Fukue, Hiwaki, Izumi, Kadusa, and Kikai). These datasets were generated using the “-p 3 -r 0.70” (without Kyushu) and the “-p 4 -r 0.70” (with Kyushu) option, respectively, to assess the genetic population structure and the history of the boundary population for the Tajima-Tango group.

The fifth was the “Local” dataset, including two Kyushu deme samples: one was Umeshiro (US), which was sampled in the northern part of the Saga prefecture, and the other was Ogi (SO), which was sampled in the southern part of the Saga prefecture. These samples were used to estimate the time of Honshu’s population divergence from Kyushu to infer the timing of the “Out of Northern Kyushu” event (Katsumura *et al*. 2012).

Note that we define the terms “deme samples” and “non-deme samples” as “samples from the local-wild population” and “samples from the wild lab stocks”, respectively (see details in Katsumura *et al*. 2009).

Against those datasets (except “PopStat” and “HZ-1”), the SNPs with strong linkage disequilibrium were randomly removed, one from each pair of SNPs with r^2^ > 0.2, using Plink1.9 (Chang *et al*. 2015) and the “--indep-pairwise 12.5 5 0.2 --autosome-num 24” option. These values were set based on a medaka population genomics study (Spivakov *et al*. 2014). Finally, the “Global”, “HZ-2” and “Local” datasets contained 8,361 SNPs out of 13,177 SNPs, 1,014 SNPs out of 2,661 SNPs and 698 SNPs out of 2,246 SNPs, respectively.

### Genetic clustering analysis

To obtain a genetic overview of the relationship of medaka geographic populations, we performed a principal component analysis (PCA) of the “Global” dataset as implemented in the *SNPRelate* program (Zheng *et al*. 2012) in R version 3.2.2. Additionally, to examine the genetic relationships between medakas in the Japanese archipelago, we used the subdataset without the E/W.KOR and Chinese populations, which included 7,126 SNPs. We also performed a model-based genetic clustering analysis using *ADMIXTURE v1.23* (Alexander *et al*. 2009) to estimate the proportions of ancestral medaka populations. We ran 50 replicates with random seeds for the number of clusters (K) from 1 to 9 and calculated the mean of the lowest fivefold cross-validation errors for each K (Jeong *et al*. 2016). Values of K = 4, 5, and 6 showed the first-, second- and third-lowest fivefold cross-validation errors, respectively (supplementary fig. 2). The results of the genetic clustering analyses were visualized by a ggplot2 package in R (Wickham 2011).

### Reconstructing the phylogenetic tree using the maximum likelihood method

We considered an individual as representative of a population and generated the individual sequences using the *population* program with the “-p 58 -r 1.00 --phylip_all” option in the *Stacks* pipeline. The dataset included 4,638 partitions and 217,986 bp of nucleotide sequences to compensate for the loci that could not be sequenced by invariable sites across all samples and “N” at the variable site. The dataset including 4,638 partitions and 217,986 bp nucleotide sequences was analyzed via the *IQ-TREE* program (Nguyen *et al*. 2015) to reconstruct a maximum likelihood tree with model selection for each partition (Chernomor *et al*. 2016) and 1,000 SH-aLRT/ultrafast-bootstrappings (Guindon *et al*. 2010; Minh *et al*. 2013). Then, we used the *FigTree* program (http://tree.bio.ed.ac.uk/software/figtree/) to visualize the phylogenetic tree.

### Inference of demographic parameters of the populations in and around Tajima-Tango and the time of “Out of Northern Kyushu” based on the ABC framework

Using the HZ-2 dataset, which added five Kyushu populations to 15 boundary populations of the HZ-1 data set, the population history, including the possibility that the Tajima-Tango group occurred by the admixture of N.JPN and S.JPN, was inferred from 1,014 SNPs using *DIYABC ver2.1.0* (Cornuet *et al*. 2014). We tested four evolutionary scenarios in which N.JPN and S.JPN diverged from an ancestral population at time t3: (I) Tajima-Tango originated in N.JPN: Honshu and Kyushu diverged at time t2, and then Tajima-Tango diverged from N.JPN at time t1 (Takehana *et al*. 2016); (II) Admixture of N.JPN and Honshu: Honshu and Kyushu diverged at time t2, and then Tajima-Tango occurred at time t1 by an admixture with rate r between N.JPN and Honshu (Sakaizumi 1984); (III) N.JPN originated in Tajima-Tango: Honshu and Kyushu diverged at time t2 and then N.JPN diverged from Tajima-Tango at time t1; and (IV) Honshu diverged from Kyushu and then Tajima-Tango occurred by an admixture with Honshu of rate r. For simplicity, all populations were assumed to have constant effective sizes in each lineage (i.e., no bottleneck and expansion). The same prior parameters were defined for four scenarios based on previous studies, and the prior distribution of each parameter is presented in table 2. In addition, we set the conditional constraint as follows: t3 > t2, t3 > t1 and t2 ≥ t1. In total, four million simulations were run, which provided approximately one million simulations for each scenario.

**Table 2.**
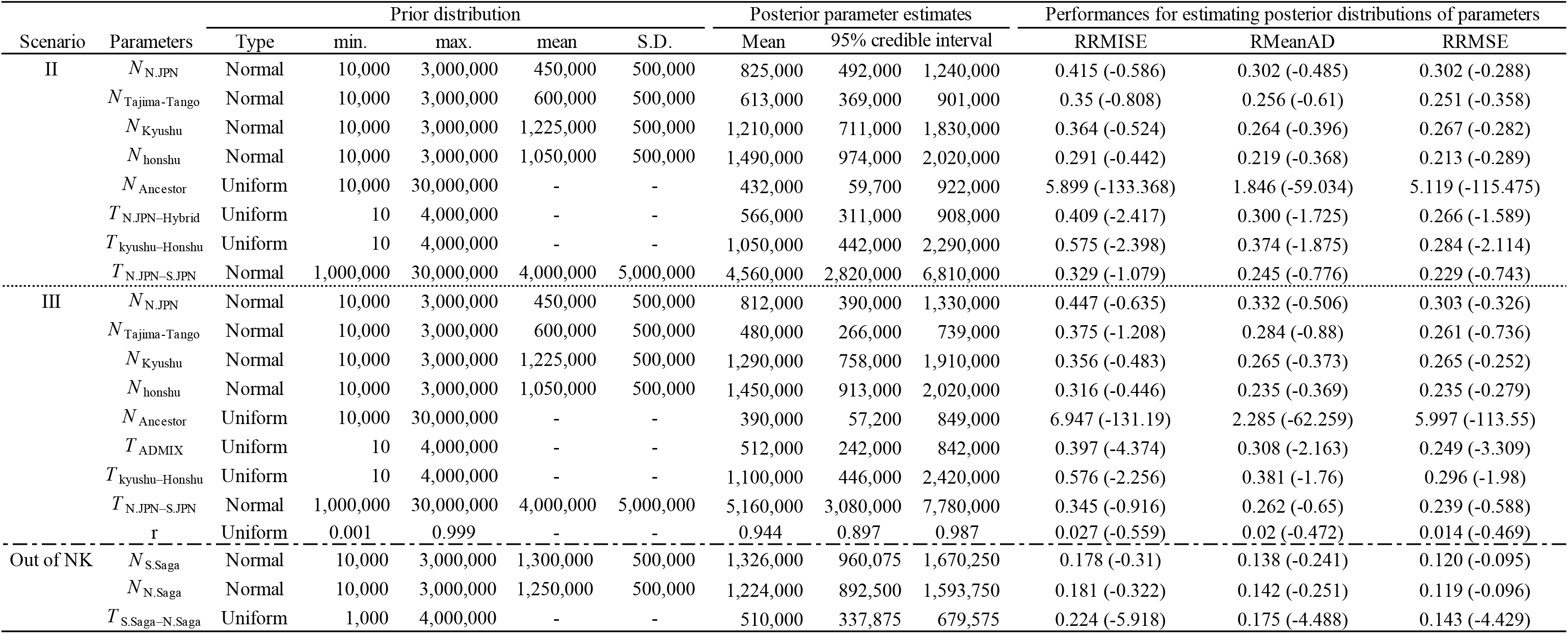
Demographic parameters estimated by DIYABC under three scenarios. Performance of each estimation was evaluated by RRMISE (the square Root of the Relative Mean Integrated Square Error), RMeanAD (the Relative Mean Absolute Deviation) and RRMSE (the square Root of teh Relative Mean Square Error), which are output from DIYABC option “Compute bias and mean square error.” The numbers with or without parentheses in the culunms “Performances for estimating posterior distributions of parameters” are those of the statistics computed from the prior or posterior distribution of parameters, respectively.

The summary statistics for all SNP loci included the proportion of loci with null gene diversity, mean gene diversity across polymorphic loci, proportion of loci with null Nei’s distance between the two samples, variance across loci of non-null Nei’s distances between two samples, proportion of loci with null admixture estimates, mean across loci of non-null admixture estimates, variance across loci of non-null admixture estimates and mean across all locus admixture estimates. The 1% simulated datasets closest to the observed dataset were used to estimate the posterior parameter distributions through a weighted local linear regression procedure (Beaumont *et al*. 2002). Scenarios were compared by estimating their posterior probabilities using the direct estimation and logistic regression methods implemented in *DIYABC* (Cornuet *et al*. 2014). We also estimated the time for “Out of Northern Kyushu” using the Local dataset that included 698 SNPs from *DIYABC*. We inferred the following time-based simple evolutionary scenario: N.Saga diverged from S.Saga at time t because this divergence event was consistent with the “Out of Northern Kyushu” event (Katsumura *et al*. 2012). In this case, all populations were assumed to have unchangeable effective sizes in each lineage.

The prior distribution of each parameter is presented in table 2, and the 1% simulated datasets closest to the observed dataset were used to estimate the posterior parameter distributions. We evaluated the accuracy of the demographic parameter estimation by calculating accuracy indicators (*Bias and mean square error* option) in *DIYABC* and accepted the parameters that were non-scaled and scaled by the mean effective population size for the HZ-2 and Local datasets, respectively. Using the infinite-sites model in the Wright-Fisher populations of a constant size, a rough value of *N*_e_ was estimated using the relationship π = 4*N*_e_μ (Tajima 1983), where π is genome-wide nucleotide diversity and μ is the mutation rate per site per generation (Osada 2015). We calculated *N*_e_ using the above formula with the following values: μ = 10^−9^/site/generation; generation = 1 year, which resulted in a mean effective population size from all presented demes in northern Kyushu of 1,275,000.

### Reconstruction of the phylogenetic tree for the partial mitochondrial DNA sequence

To examine the mitochondrial introgression in the tested samples, we generated and analyzed the mitochondrial DNA (mtDNA) partial sequences from the GBS-read data. We mapped the reads to the complete mtDNA sequence of “Clade C” (Matsuda *et al*. 1997; Takehana *et al*. 2003), which diverged genetically from S.JPN and N.JPN, to eliminate the mapping bias that occurs when genetically distant sequences are used as a reference. The genomic DNA of “Clade C” was extracted in Katsumura *et al*. 2009. The complete mtDNA sequence was determined as follows. Using PCR, five DNA fragments of mitochondrial DNA were amplified: fragment 1—4,556 bp; fragment 2—4,527 bp; fragment 3-1—4,589 bp; fragment 3-2—4,504 bp; and fragment 4—4,546 bp (supplementary fig. 3). Supplementary table 1 describes the primers, which were designed based on the inbred strain Hd-rR complete mtDNA sequence (accession number: AP008938). Approximately 20 ng of the genomic DNA was used as a template for the PCR assay in a 50 μl solution containing dNTP at 0.2 mM, 0.2 μM of each of primer, 0.75 U of EX Taq polymerase HS (TaKaRa Shuzo Co.), and the reaction buffer attached to the polymerase. The reactions were conducted in a TaKaRa PCR Thermal Cycler Dice (TaKaRa Shuzo Co.) using the following protocol: an initial denaturing step at 95°C for 2 min, 40 cycles of denaturation at 95°C for 30 sec, annealing at 60°C for 30 sec, extension at 72°C for 300 sec, and a final extension step at 72°C for 5 min. The PCR products were diluted 20-fold and used as templates in the sequencing reaction (following the commercial protocol) with thirty-three primers (supplementary table 1, supplementary fig. 3) and then analyzed in an ABI 3500xL Genetic Analyzer (Life Technologies). The complete mtDNA sequence of “Clade C” was reconstructed using *SeqMan Pro 10.1.2.20* (DNASTAR) and deposited into the international DNA database DDBJ/EMBL/GenBank (accession number: LC335803).

The “Clade C” complete mtDNA sequence was used to align the reads using *BWA backtrack 0.7.12-r1039* (Li and Durbin 2009) using the “-n 5” option. After the mapping process, to remove the multi-mapped reads, we used *Samtools v1.2* (Li *et al*. 2009) using the “-Sq 20” option. Then, we used *Stacks* with the “-m 3” option for minimum stack depth (Catchen *et al*. 2011; 2013) and then obtained the 322 bp nucleotide sequences. The loci where the sequence was missing were filled in with “N”. In addition, the nucleotide position showing the multi-allelic state was replaced by “N”. Phylogenetic trees were constructed using the neighbor-joining (NJ) method (Saitou and Nei 1987) with the program MEGA5 (Tamura *et al*. 2011). The evolutionary distances were calculated using the Jukes-Cantor method (Jukes and Cantor 1969). The analysis also involved 13 nucleotide sequences from the DNA database (See supplementary fig. 4). All ambiguous positions were removed for each sequence pair. The reliability of the tree was evaluated using 1000 bootstrap replicates (Felsenstein 1985).

## Results

### Autosomal genetic diversity of five groups of *Oryzias latipes*

To assess the genetic diversity based on autosomal SNPs within known mitochondrial and hybridization groups—the northern Japanese (N.JPN), southern Japanese (S.JPN), eastern Korean (E.KOR) western Korean/Chinese (W.KOR) and Tajima-Tango groups—their population-genetic summary statistics were calculated using the “PopStat” dataset that included the alignment regions across all groups and contained approximately 45 kb of sequences and 2,453 SNPs (table 1). All groups showed major allele frequencies and heterozygosities of 0.940–0.983 and 0.023–0.054, respectively, indicating many rare alleles and homozygous sites in our wild lab-stocks. W.KOR showed the highest nucleotide diversity (0.0053 ± 0.0003) in all lab-stocks originated from East Asia. S.JPN showed the highest nucleotide diversity (0.0036 ± 0.0001) and N.JPN the lowest in the Japanese archipelago (table 1), which is consistent with the diversity based on mtDNA (Katsumura *et al*. 2009).

### Genetic clustering based on autosomal SNPs inferring medaka population structure

To reveal the medaka population structure based on the autosomal SNPs, we performed genetic clustering analyses using the “Global” dataset. First, we investigated the genetic relationship by a principle component analysis (PCA) using 8,361 SNPs on 24 autosomes. The PCA showed that the wild lab-stocks were divided into two clusters, N.JPN/Tajima-Tango and S.JPN and others from E/W.KOR, which is similar to the grouping based on the mtDNA data (Takehana *et al*. 2003; Katsumura *et al*. 2009)(fig. 2a, supplementary fig. 5). E/W.KOR individuals were plotted dispersedly compared with other clusters, and Yongcheon (E.KOR) was close to N.JPN/Tajima-Tango. As PC3 was divided into E.KOR and W.KOR, the autosomes in these mtDNA groups were differentiated into each other.

**FIG. 2:**
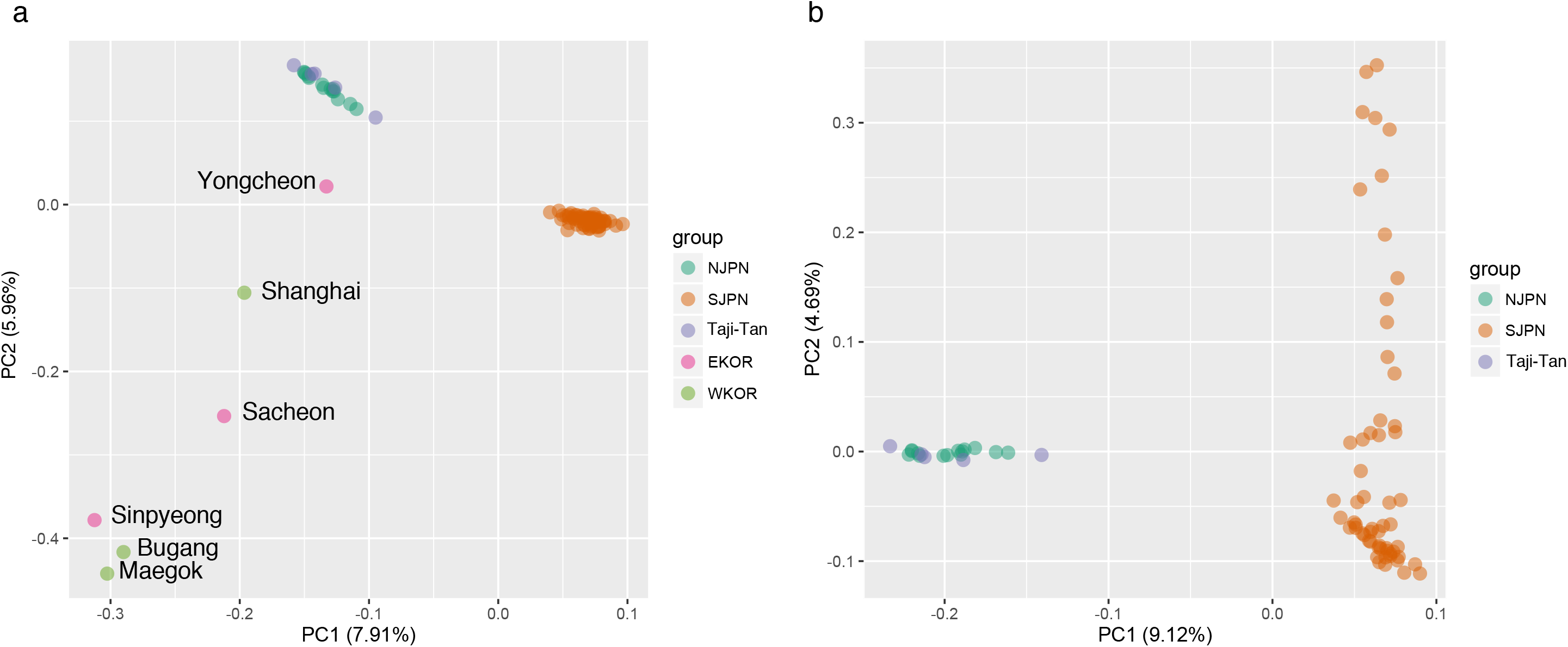
Results of principle component analysis (PCA) using SNPs in East Asia (a) and the Japanese archipelago (b). Each plot shows PC1 versus PC2. The population names of each point are described in fig. S6.

Although Tajima-Tango was considered to be a hybridization group between N.JPN and S.JPN (Sakaizumi 1984), all individuals from Tajima-Tango were plotted with those from N.JPN as a cluster (fig. 2a). Next, for a fine-scale mapping of Japanese medaka, we performed the PCA based on 7,126 SNPs, excluding E/W.KOR individuals (fig. 2b). The PCA showed that the Kyushu populations (Kudamatsu, Ogi, Izumi, Hiwaki, Kikai, Nago, Gushikami, Hisayama, Umeshiro, Ashibe, Arita, Kusu and Nobeoka) in S.JPN were dispersed along the PC2 axis (supplementary fig. 6), while that Tajima-Tango overlapped with N.JPN again, even though we used up to PC5 (supplementary fig. 5). This result indicated that Tajima-Tango was not considerably differentiated from N.JPN on autosomes, suggesting that S.JPN was differentiated within the groups and the genetic diversity was especially high among the Kyushu populations (fig. 2b).

To examine the phylogenetic relationship among 83 populations, we constructed a maximum likelihood (ML) tree based on 4,638 short fragment sequences (comparative nucleotide sequence length: 217,986 bp) using *IQ-TREE*. The maximum likelihood tree showed relatively high bootstrap values on each branch and almost the same topology as previous trees based on mtDNA (Takehana *et al*. 2003; Katsumura *et al*. 2009), except for Yongcheon (fig. 3 and supplementary fig. 7; see also Discussion). Medakas from the Korean peninsula were genetically close to each other in both groups, whereas those from Shanghai, which have been classified into W.KOR based on mtDNA studies, diverged from the other individuals from W.KOR. The medakas in the Japanese archipelago were divided into two major clusters similar to mtDNA. One was the N.JPN/Tajima-Tango, and the other was S.JPN. The N.JPN/Tajima-Tango cluster was further divided into two submajor clusters with a 100% bootstrapping value in the ML tree based on concatenated sequences obtained from GBS, although N.JPN and Tajima-Tango overlapped in PCA, which reduced SNP information, i.e., N.JPN and Tajima-Tango diverged on the whole autosomal sequenced regions. S.JPN was also further divided into two sub-major clusters: Kyushu-only and Kyushu & others, which included 14 Kyushu populations, represented in fig. 3 by an open and a closed red circle, respectively. While the Kyushu-only cluster diverged into the northern and the southern Kyushu, the Kyushu & others cluster diverged to the Pacific side of eastern and the northern Honshu (main-island Japan), which included several clusters supported by high bootstrapping value (SH-aLRT ≥ 80% and UFboot ≥ 95%, supplementary fig7). Finally, we characterized the S.JPN ancestry in the context of East Asian genetic diversity of medaka by performing an *ADMIXTURE* analysis, which is a model-based unsupervised genetic clustering method. With the optimal number of ancestral components (K = 4), S.JPN medakas were assigned to two distinct ancestries (fig. 3 and supplementary fig. 2). In suboptimal runs with more ancestral components (K = 5, 6), only S.JPN in Honshu (main-island Japan) was assigned to the other ancestral components found in the northern Kyushu populations. This analysis indicated that genetic diversities in the northern Kyushu populations were the highest among S.JPN. Thus, the genetic clustering analyses based on genome-wide SNP data strongly suggested that S.JPN spread from northern Kyushu to Honshu.

**FIG. 3:**
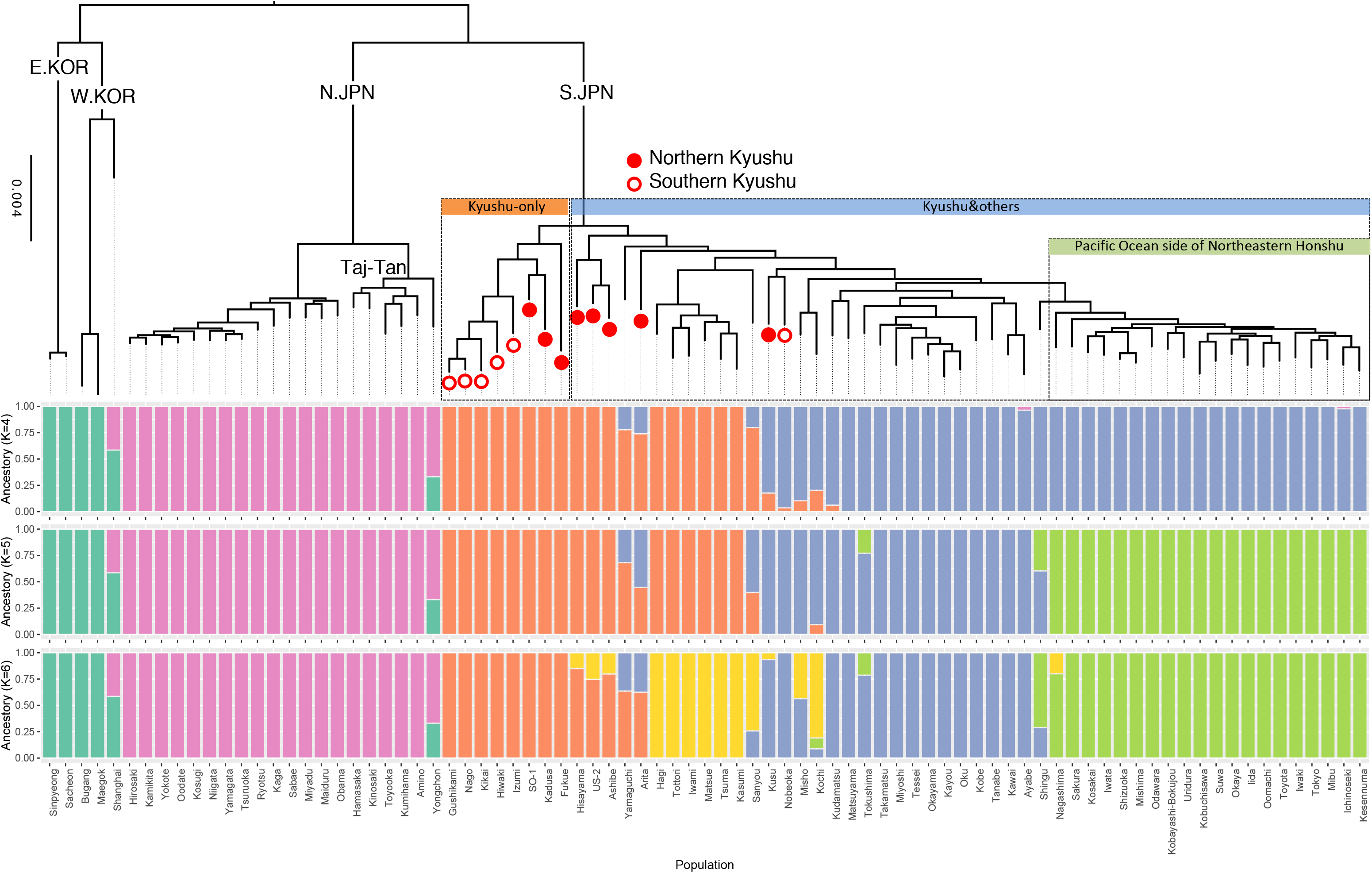
Phylogenetic tree using the maximum likelihood method and an ancestry barplot with ADMIXTURE analysis. Red closed and open circles represent the northern and southern Kyushu populations, respectively. “Taj-Tan” in the tree is the abbreviation for Tajima-Tango.

### Boundary population genomes similar to that of the northern Japanese group

To explore hybridization signatures on autosomes, we examined allele-sharing between boundary populations and surrounding populations using the “HZ-1” dataset (see Materials and Methods). We classified the fixed alleles between the groups into three states: shared by N.JPN and Tajima-Tango, shared by S.JPN and Tajima-Tango, and shared by N.JPN and S.JPN. Then, we summarized each state’s total numbers, as shown in table in fig. 4. We found 1,380 out of 4,661 SNPs that were common alleles between two of the three groups (fig. 4). Regarding those alleles, the majority (81.4%) were common alleles between Tajima-Tango and N.JPN, which was a much higher frequency than that between Tajima-Tango and S.JPN (11.2%). The rest of them were common alleles between N.JPN and S.JPN (7.5%), i.e., specific alleles in Tajima-Tango. These proportions are near those of a previous study based on 96 genomic regions (4 loci for each chromosome) (Takehana *et al*. 2016). Although the regions with neighboring common alleles between Tajima-Tango and S.JPN were observed for certain chromosomes, alleles on the Tajima-Tango genome were mostly shared with N.JPN, suggesting that Tajima-Tango is a subgroup of N.JPN with a short divergence time.

**FIG. 4:**
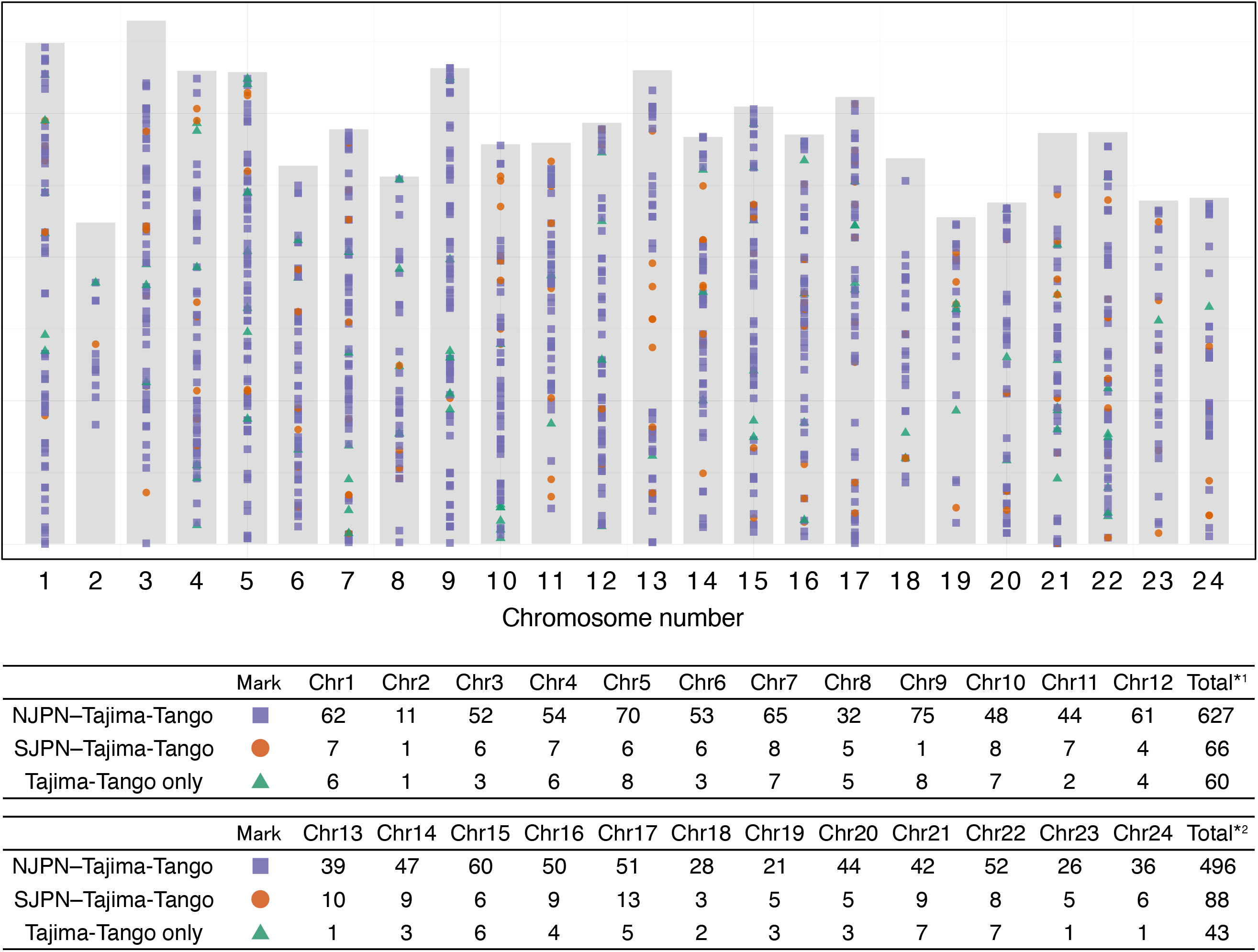
Shared allele distribution among boundary populations. The gray rectangle represents the total length of each medaka chromosome. The table in the figure shows the number of alleles shared between the groups observed on each chromosome. Mark represents the state whose group shares that allele. *1 and *2 are the total number from chromosomes 1 to 12 and from chromosomes 13 to 24, respectively.

To investigate the mitochondrial introgression in boundary populations, we reconstructed the phylogenetic tree based on partial mtDNA sequences generated by mapping short reads from the GBS of the complete mitochondrial genome (supplementary fig. 4). The mtDNA sequences from Toyooka and Kinosaki in Tajima-Tango were clustered and closely related to those from Kaga, which was a root population in N.JPN. Considering that Tajima-Tango was a subgroup of N.JPN, which we inferred from the ML tree based on autosomal sequences, these phylogenetic positions on the mtDNA tree would have also reflected an evolutionary history in which N.JPN diverged from the Tajima-Tango group. We confirmed that Amino, Kumihama and Hamasaka in Tajima-Tango had the mitochondrial genomes of phylogenetically distant populations classified into S.JPN (supplementary fig. 4). Additionally, the mitochondrial genome of Ayabe in S.JPN was classified into N.JPN but diverged from Tajima-Tango. These results suggest that mitochondrial genome introgressions occurred reciprocally, meaning that they occurred not only from S.JPN to Tajima-Tango but also from N.JPN to S.JPN.

### Inferring medaka demographic parameters

To infer the demographic parameters, effective population size (*N*_e_), time (T), and proportion of the admix (r), we analyzed the “HZ-2” dataset based on the coalescent theory using an approximate Bayesian computation (ABC) framework. We performed the model selection to identify the best explanation scenario for the observed data from the four scenarios; (I) Tajima-Tango originated in N.JPN, (II) Admixture of N.JPN and Honshu, (III) N.JPN originated in Tajima-Tango, and (IV) Admixture of Tajima-Tango and Honshu (fig. 5a). *DIYABC* has two different approaches (directional and logistic) for model selection. Based on each criterion, scenario III (Honshu diverged from Kyushu and then N.JPN diverged from the Tajima-Tango group without admixture with Honshu) and scenario IV (Honshu diverged from Kyushu and then Tajima-Tango occurred by an admixture with Honshu at r rate) were supported by the directional (posterior probability: 0.4720, 95% CI: 0.0344–0.9096) and logistic (posterior probability: 0.4477, 95% CI: 0.4313–0.4641) approaches, respectively (supplementary fig. 8). Our data could not distinguish between scenarios III and IV (see the Discussion section).

**FIG. 5:**
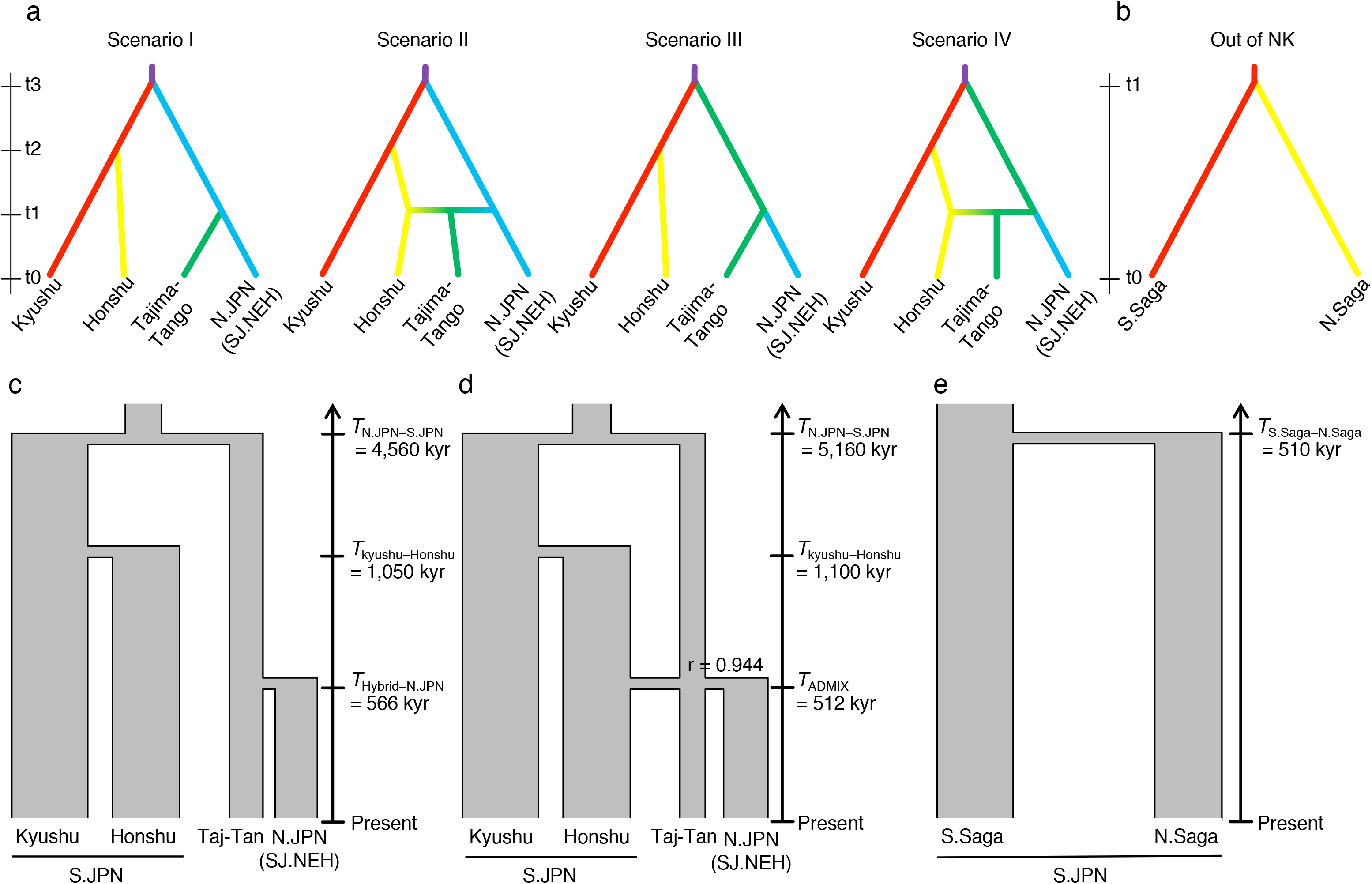
Scenarios for estimation of the demographic parameters. Possible scenarios for the population history of the Tajima-Tango group (a) and the “Out of northern Kyushu (NK)” event (b) are shown. The TMRCAs estimated using ABC are in scenario3 (c), scenario4 (d) and Out of NK (e).

The posterior parameter estimates of scenarios III and IV shown in table 2 and figs. 5c and 5d were not scaled because various measures of accuracy (RRMISE, RMeanAD, and RRMSE in the *DIYABC* output) indicated that non-scaled parameters fit the observed data better than scaled parameters Additionally, although we inferred the time to the most common ancestor at three divergence events (fig. 5), these times might be overestimated because the dataset for this estimation consisted of non-deme samples, i.e., polymorphisms within populations were underestimated, and the random-mating assumption could not be satisfied. Therefore, we estimated the timing of the “Out of Northern Kyushu” event using the “Local” dataset composed of the two deme samples (S.Saga and N.Saga), which were split at first on the S.JPN lineage (fig. 3). Based on a simple hypothesis, constant population size and no migration, we obtained the estimated time (510,000 years ago, 95% CI: 337,875–679,575) for the ancestral divergence of the two deme samples (table 2 and fig. 5e), which was calculated from a scaled parameter by the mean effective population size because various accuracy measures indicated that scaled parameters fit the observed data better than non-scaled parameters. Thus, our population-genetic estimate using ABC suggests that medakas diverged and dispersed throughout Pacific-side Japan approximately 510 kyr ago.

## Discussion

### Redefined subgroups in Japanese archipelago based on the population structure inferred from genome-wide single-nucleotide polymorphisms

*Oryzias latipes* can be divided into five groups by mtDNA sequences and allozymic electrophoresis patterns (Sakaizumi 1984; Takehana *et al*. 2003). In this study, based on autosomal SNPs, the genetic clustering analysis showed that “K = 4” was the most supportive because it presented the lowest fivefold cross-validation error, indicating that N.JPN and S.JPN were divided into three ancestral clusters. When the K values increased, only S.JPN divided into more subgroups, which suggests that the S.JPN group was composed of more divergent groups than the other groups. Considering together with the results of the ML tree analysis, it is possible to redefine subgroups composed of each major group for our wild lab-stocks originated from the Japanese archipelago as follows. First, Tajima-Tango, which had been considered a hybridization group, should be included under the N.JPN group because it shows almost the same ancestral component as N.JPN. Then, the group called N.JPN should be assigned to a subgroup, for which we propose the name “Sea of Japan side of Northeastern Honshu (SJ.NEH).” These two subgroups, Tajima-Tango and SJ.NEH, compose the N.JPN group (supplementary fig. 7). Second, S.JPN can be divided into several subgroups, San-in, San-yo/Shikoku/Kinki and the Pacific Ocean side of Northeastern Honshu (PO.NEH), because the Kyushu & others cluster was subdivided into three sub-clusters composed of geographically neighboring populations. Adding the Kyushu subgroup, S.JPN is composed of four subgroups. Thus, medaka in Japanese archipelago could also be composed of the six distinct subgroups based on autosomal genetic diversity (supplementary fig. 7).

S.JPN can be divided into finer subgroups based on the mitochondrial genome (Takehana *et al*. 2003), likely because the effective population size of mtDNA is one quarter that of a nuclear gene. This causes the intermingled branch pattern of the mtDNA tree, which is not associated with geographic distance. Although this pattern has been observed our wild lab-stocks in a previous study based on mtDNA (Katsumura *et al*. 2009), it has been interpreted as artificial migrations accompanied by recent human activities (Takehana *et al*. 2003). If the branching pattern was formed by human activities, various ancestral components should appear independent of geographic relatedness in the result of the *ADMIXTURE* analysis based on autosomal SNPs. However, no signal of recent migration was found in the *ADMIXTURE* result. Rather, the autosomal SNP data support another hypothetical scenario in which the mtDNA tree topology reflects ancestral polymorphisms only and their local fixation is caused by a small effective population size (Katsumura *et al*. 2009). This result indicates that arguing the geographical origins of medaka based only on mtDNA may lead to false conclusions.

From our data, it may be difficult to accurately evaluate the genetic diversities in E/W.KOR groups because the number of populations examined in each group was small. Therefore, though care must be taken in the interpretation, W.KOR showed the highest nucleotide diversity and E.KOR the second highest among the four major mitochondrial groups (table 1). The W.KOR group included the Chinese medaka in Shanghai, which could have elevated its value, while the latter group did not include any geographically distant populations. The clustering analysis showed that Shanghai from W.KOR and Yongcheon from E.KOR had an ancestry component from N.JPN (fig.3). This suggests two possibilities: one is that the N.JPN ancestor was derived from the E/W.KOR ancestor, and the other is that contamination occurred through the maintained wild lab stocks. To investigate these possibilities and the origin of *O. latipes*, a population-based genome wide analysis must be conducted to increase the population numbers of E/W.KOR and include the sister species *O. curvinotus* and *O. luzonensis*.

### Reconstruction of the medaka population history in the Japanese archipelago

Our genome-wide analysis shows that medakas in N.JPN and S.JPN are deeply divergent and dispersed over the Japanese archipelago from different locations at different times. In particular, our ABC analysis indicates that SJ.NEH originated in and diverged from Tajima-Tango. This inference about the history of N.JPN after divergence from S.JPN is not consistent with previous inferences from the allozyme, mtDNA and limited autosomal SNP analyses, which have suggested that Tajima-Tango is a hybridization group between PO.NEH and S.JPN, or Tajima-Tango is a sub-group derived from PO.NEH. Our GBS data show that (i) the nucleotide diversity in Tajima-Tango is higher than that in SJ.NEH (table 1), (ii) S.JPN is genetically more closely related to Tajima-Tango than to SJ.NEH based on the allele-sharing rates (fig. 4), and (iii) the Tajima-Tango branch is the root in the N.JPN clade of the phylogenetic tree based on partial mtDNA sequences (supplementary fig. 4), which is also shown by the whole mitochondrial genome analysis (Hirayama *et al*. 2010). These data suggest that Tajima-Tango forms an outgroup to all present-day SJ.NEH and it spread along the Sea of Japan side. Furthermore, the ABC framework’s estimation supports our scenario, although the analysis does not statistically distinguish between scenario III and scenario IV. Even if the admixture occurred, the inferred ratio of the admixture is too low (fig. 5d).

Our findings strongly support the “Out of northern Kyushu” model of S.JPN proposed in Katsumura et al. (2012) and have revealed the dispersal route of SJ.NEH/Tajima-Tango in N.JPN. The genetic clustering analyses and phylogenetic tree based on the GBS data elucidate medaka history better than previous mitochondrial DNA analyses. In particular, the *ADMIXTURE* analysis shows that the two ancestry components (yellow and blue in fig. 3) were observed in northern Kyushu, suggesting that S.JPN dispersed in three different directions after dispersing out of Northern Kyushu. Geographically, Shikoku, Sanyo, and Kinki are separated by sea, but medakas in the three local lands share the same ancestral component (blue in fig.6; see also supplementary fig. 1 for geographic information). Medaka is a freshwater fish, but survival and reproduction are possible even in seawater (Inoue and Takei 2002). Because most of the rivers in the Japanese archipelago are steep and short, the rivers tend to flood after heavy rain. Although medakas are highly likely to drift to the sea on each occasion, medakas can have survived even in the sea and may have returned to the river because of their saltwater tolerance. Thus, the possibility of moving from river to river through the sea cannot be ignored. The results of this study suggest a history in which medakas migrated throughout the Japanese archipelago through the sea.

The different branching patterns of autosomal and mtDNA trees suggest that the mtDNA introgression occurred not only from S.JPN to Tajima-Tango but also from N.JPN to S.JPN. The Tajima-Tango region is surrounded by mountains (supplementary fig. 1); however, it contains the lowest watershed (sea level 95.45 m) in the Japanese archipelago.

The medaka has possibly moved in both directions across the watershed. From the above, the most plausible scenario is as follows (fig. 6). Ancestral S.JPN and N.JPN diverged first and independently reached their current habitats in the Japanese archipelago. After S.JPN, which is an ancestor of San-in, San-yo/Shikoku/Kinki and PO.NEH, dispersed from northern Kyushu approximately 510 kyr ago, SJ.NEH diverged from Tajima-Tango in N.JPN. Then, their descendant populations spread rapidly to northern Honshu on the Sea of Japan side. Meanwhile, S.JPN dispersed in as many as three different directions and then spread rapidly northeastward from the western part of Fossa Magna. In the process of dispersing across the main island of Japan, certain S.JPN populations infiltrated the Tajima-Tango region from the west and the south, and the resulting mtDNA introgression occurred independently.

**FIG. 6:**
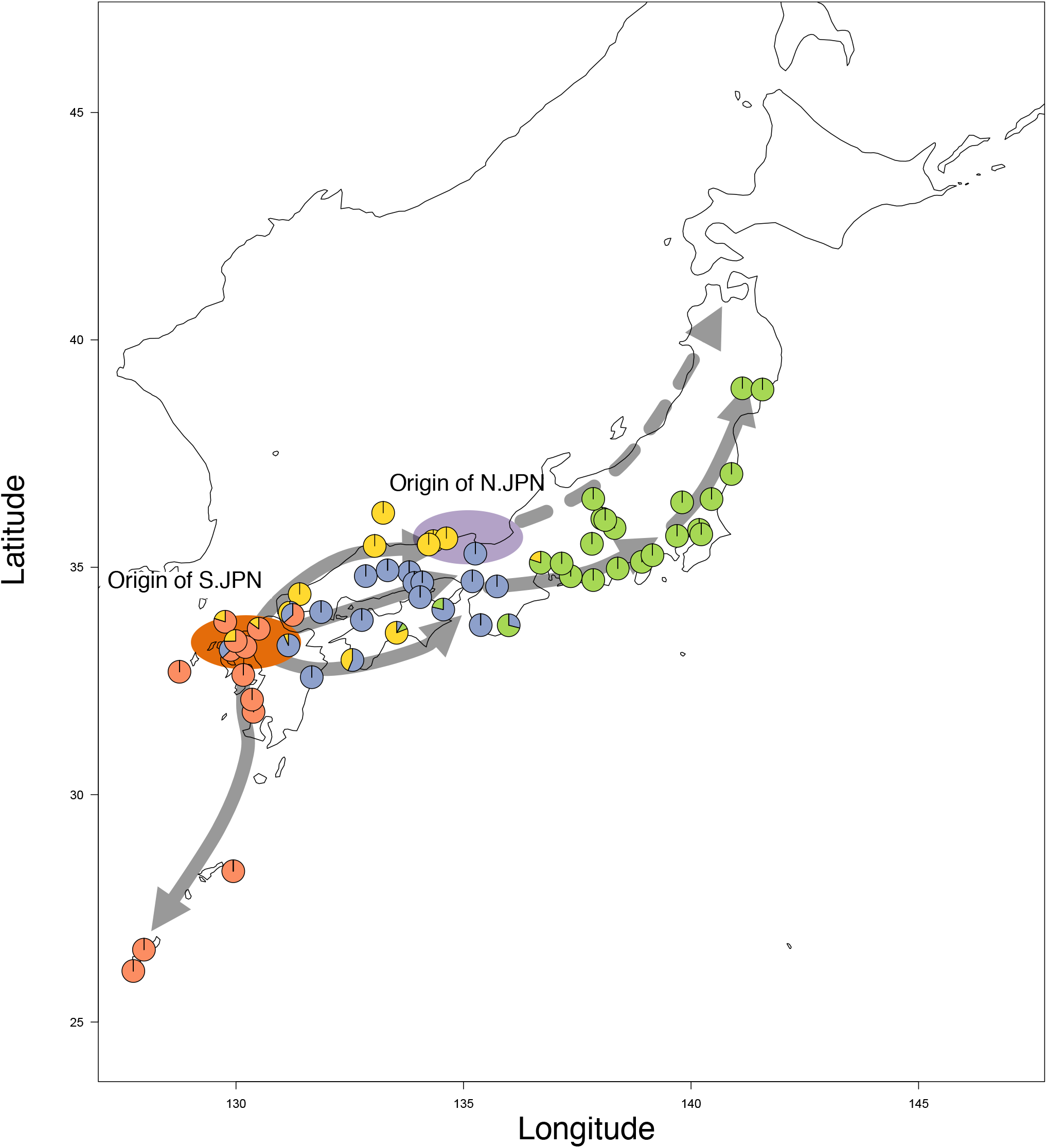
Map representing the ancestry proportions from ADMIXTURE analysis at K = 6. Solid and dashed lines represent the spreading patterns of S.JPN and N.JPN inferred by GBS data, respectively.

## Conclusion

Our genome-wide SNP analysis reconstructed the detailed population structure and reliable history of medaka that evolved in the Japanese archipelago. Since the distribution of the subgroups was highly consistent with the geographical features, several adaptive traits could have evolved in each subgroup. Furthermore, the boundary populations were not caused by a hybridization event but instead were the origin of the populations dispersed to a northeastern part of the Japanese archipelago on the Sea of Japan side. A better understanding of the population structure and history of medaka will support association studies for phenotypes and genotypes related to environmental adaptation.

## Acknowledgments

We thank Mrs. Shizuko Chiba, Mrs. Sumiko Tomizuka, Dr. Atsuko Shimada and Prof. Emeritus Akihiro Shima (the University of Tokyo) for maintaining the medaka stocks from wild populations. This study was supported in part by grants-in-aid for Young Scientists (B), no. 16K21352 to T.K., and by a grant-in-aid for Scientific Research (A), nos. 26251050, 25251046 and 17H01453 to H.O. T.K. was also supported by a grant-in-aid from JSPS Research Fellowships (16J07227).

## References

Alexander, D. H., J. Novembre, and K. Lange, 2009 Fast model-based estimation of ancestry in unrelated individuals. Genome Research 19: 1655–1664.

Andrews, K. R., J. M. Good, M. R. Miller, G. Luikart, and P. A. Hohenlohe, 2016 Harnessing the power of RADseq for ecological and evolutionary genomics. Nature Review Genetics 17: 81–92.

Ansai, S., and M. Kinoshita, 2014 Targeted mutagenesis using CRISPR/Cas system in medaka. Biology Open 3: 362–371.

Beaumont, M. A., W. Zhang, and D. J. Balding, 2002 Approximate Bayesian computation in population genetics. Genetics 162: 2025–2035.

Catchen, J., P. A. Hohenlohe, S. Bassham, A. Amores, and W. A. Cresko, 2013 Stacks: an analysis tool set for population genomics. Mol Ecol 22: 3124–3140.

Catchen, J. M., A. Amores, P. Hohenlohe, W. Cresko, and J. H. Postlethwait, 2011 Stacks: building and genotyping Loci de novo from short-read sequences. G3 (Bethesda) 1: 171–182.

Chang, C. C., C. C. Chow, L. C. Tellier, S. Vattikuti, S. M. Purcell et al., 2015 Second-generation PLINK: rising to the challenge of larger and richer datasets. Gigascience 4: 7.

Chernomor, O., A. von Haeseler, and B. Q. Minh, 2016 Terrace Aware Data Structure for Phylogenomic Inference from Supermatrices. Systematic Biology 65: 997–1008.

Cornuet, J.-M., P. Pudlo, J. Veyssier, A. Dehne-Garcia, M. Gautier et al., 2014 DIYABC v2.0: a software to make approximate Bayesian computation inferences about population history using single nucleotide polymorphism, DNA sequence and microsatellite data. Bioinformatics 30: 1187–1189.

Elshire, R. J., J. C. Glaubitz, Q. Sun, J. A. Poland, K. Kawamoto et al., 2011 A robust, simple genotyping-by-sequencing (GBS) approach for high diversity species. PLoS ONE 6: e19379.

Felsenstein, J., 1985 CONFIDENCE LIMITS ON PHYLOGENIES: AN APPROACH USING THE BOOTSTRAP. Evolution 39: 783–791.

Guindon, S., J.-F. Dufayard, V. Lefort, M. Anisimova, W. Hordijk et al., 2010 New algorithms and methods to estimate maximum-likelihood phylogenies: assessing the performance of PhyML 3.0. Systematic Biology 59: 307–321.

Hirayama, M., T. Mukai, M. Miya, Y. Murata, Y. Sekiya et al., 2010 Intraspecific variation in the mitochondrial genome among local populations of Medaka Oryzias latipes. Gene 457: 13–24.

Igarashi, K., J. Kobayashi, T. Katsumura, Y. Urushihara, K. Hida et al., 2017 An Approach to Elucidate NBS1 Function in DNA Repair Using Frequent Nonsynonymous Polymorphism in Wild Medaka (Oryzias latipes) Populations (S. D. Fugmann, Ed.). PLoS ONE 12: e0170006–19.

Inoue, K., and Y. Takei, 2002 Diverse adaptability in oryzias species to high environmental salinity. Zool. Sci. 19: 727–734.

Jeong, C., S. Nakagome, and A. Di Rienzo, 2016 Deep History of East Asian Populations Revealed Through Genetic Analysis of the Ainu. Genetics 202: 261–272.

Jukes, T. H., and C. R. Cantor, 1969 Evolution of protein molecules. Mammalian protein metabolism 3: 132.

Kasahara, M., K. Naruse, S. Sasaki, Y. Nakatani, W. Qu et al., 2007 The medaka draft genome and insights into vertebrate genome evolution. Nature 447: 714–719.

Katsumura, T., S. Oda, S. Mano, N. Suguro, K. Watanabe et al., 2009 Genetic differentiation among local populations of medaka fish (Oryzias latipes) evaluated through grid- and deme-based sampling. Gene 443: 170–177.

Katsumura, T., S. Oda, S. Nakagome, T. Hanihara, H. Kataoka et al., 2014 Natural allelic variations of xenobiotic-metabolizing enzymes affect sexual dimorphism in Oryzias latipes. Proceedings of the Royal Society B: Biological Sciences 281:.

Katsumura, T., S. Oda, K. Tsukamoto, T. Yamashita, M. Aso et al., 2012 A population genetic study on the relationship between medaka fish and the spread of wet-rice cultivation across the Japanese archipelago. Anthropol. Sci.

Li, H., and R. Durbin, 2009 Fast and accurate short read alignment with Burrows-Wheeler transform. Bioinformatics 25: 1754–1760.

Li, H., B. Handsaker, A. Wysoker, T. Fennell, J. Ruan et al., 2009 The Sequence Alignment/Map format and SAMtools. Bioinformatics 25: 2078–2079.

Matsuda, M., H. Yonekawa, S. Hamaguchi, and M. Sakaizumi, 1997 Geographic variation and diversity in the mitochondrial DNA of the medaka, Oryzias latipes, as determined by restriction endonuclease analysis. Zool. Sci. 14: 517–526.

Minh, B. Q., M. A. T. Nguyen, and A. von Haeseler, 2013 Ultrafast approximation for phylogenetic bootstrap. Molecular Biology and Evolution 30: 1188–1195.

Narum, S. R., C. A. Buerkle, J. W. Davey, M. R. Miller, and P. A. Hohenlohe, 2013 Genotyping-by-sequencing in ecological and conservation genomics. Mol Ecol 22: 28412847.

Nguyen, L.-T., H. A. Schmidt, A. von Haeseler, and B. Q. Minh, 2015 IQ-TREE: a fast and effective stochastic algorithm for estimating maximum-likelihood phylogenies. Molecular Biology and Evolution 32: 268–274.

Oota, H., and H. Mitani, 2011 Human Population Genetics Meets Medaka, pp. 339–350 in Medaka: A Model for Organogenesis, Human Disease, and Evolution, edited by K. Naruse, M. Tanaka, and H. Takeda. Medaka: A Model for Organogenesis, Human Disease, and Evolution, Springer Japan, Tokyo.

Osada, N., 2015 Genetic diversity in humans and non-human primates and its evolutionary consequences. Genes Genet. Syst. 90: 133–145.

Poland, J. A., P. J. Brown, M. E. Sorrells, and J.-L. Jannink, 2012 Development of High-Density Genetic Maps for Barley and Wheat Using a Novel Two-Enzyme Genotyping-by-Sequencing Approach (T. Yin, Ed.). PLoS ONE 7: e32253–8.

Saitou, N., and M. Nei, 1987 The neighbor-joining method: a new method for reconstructing phylogenetic trees. Molecular Biology and Evolution 4: 406–425.

Sakaizumi, M., 1986 Genetic divergence in wild populations of Medaka, Oryzias latipes (Pisces: Oryziatidae) from Japan and China. Genetica.

Sakaizumi, M., 1984 Rigid Isolation between the Northern Population and the Southern Population of the Medaka, Oryzias latipes (Genetics). Zool. Sci. 1: 795–800.

Sakaizumi, M., N. Egami, and K. Moriwaki, 1980 Allozymic Variation in Wild Populations of the Fish, Oryzias latipes. Proc. Jpn. Acad., Ser. B 56: 448–451.

Sakaizumi, M., K. Moriwaki, and N. Egami, 1983 Allozymic variation and regional differentiation in wild populations of the fish Oryzias latipes. Copeia 1983: 311.

Shima, A., A. Shimada, M. Sakaizumi, and N. Egami, 1985 First listing of wild stocks of the medaka Oryzias latipes currently kept by the Zoological Institute, Faculty of Science, University of Tokyo. Journal of the Faculty of Science, Imperial University of Tokyo Sect IV, Zoology 16: 27–35.

Shimmura, T., T. Nakayama, A. Shinomiya, S. Fukamachi, M. Yasugi et al., 2018 Dynamic plasticity in phototransduction regulates seasonal changes in color perception. Nature Communications 1–7.

Spivakov, M., T. O. Auer, R. Peravali, I. Dunham, D. Dolle et al., 2014 Genomic and phenotypic characterization of a wild medaka population: towards the establishment of an isogenic population genetic resource in fish. 4: 433–445.

Tajima, F., 1983 Evolutionary relationship of DNA sequences in finite populations. Genetics 105:437–460.

Takehana, Y., N. Nagai, M. Matsuda, K. Tsuchiya, and M. Sakaizumi, 2003 Geographic variation and diversity of the cytochrome b gene in Japanese wild populations of medaka, Oryzias latipes. Zool. Sci. 20: 1279–1291.

Takehana, Y., M. Sakai, T. Narita, T. Sato, K. Naruse et al., 2016 Origin of Boundary Populations in Medaka (Oryzias latipes Species Complex). Zool. Sci. 33: 125–131.

Tamura, K., D. Peterson, N. Peterson, G. Stecher, M. Nei et al., 2011 MEGA5: Molecular Evolutionary Genetics Analysis Using Maximum Likelihood, Evolutionary Distance, and Maximum Parsimony Methods. Molecular Biology and Evolution 28: 2731–2739.

Watanabe-Asaka, T., Y. Sekiya, H. Wada, T. Yasuda, I. Okubo et al., 2014 Regular heartbeat rhythm at the heartbeat initiation stage is essential for normal cardiogenesis at low temperature. BMC Dev. Biol. 14:.

Wickham, H., 2011 ggplot2. Wiley Interdisciplinary Reviews: Computational Statistics 3: 180–185.

Zheng, X., D. Levine, J. Shen, S. M. Gogarten, C. Laurie et al., 2012 A high-performance computing toolset for relatedness and principal component analysis of SNP data. Bioinformatics 28: 3326–3328.

